# The Endosomal pH Regulator NHE6 is Required for Insulin-Stimulated Glucose Uptake in Adipocytes

**DOI:** 10.1101/2023.07.30.551163

**Authors:** Allatah X. Mekile, Monish Ram Makena, Ruby Gupta, Cory J. White, Ray Wesley Bowman, Rachana L. Patnayak, Damaris N. Lorenzo, Rajini Rao

**Affiliations:** Department of Physiology, The Johns Hopkins University School of Medicine, Baltimore, MD, USA; Department of Biological Chemistry, The Johns Hopkins University School of Medicine, Baltimore, MD, USA.; Department of Cell and Developmental Biology, University of Pennsylvania Perelman School of Medicine, Philadelphia, PA

**Keywords:** Na^+^/H^+^ exchanger, GLUT4, diabetes, metformin, insulin

## Abstract

**Objective:** Trafficking of <u>GLU</u>cose <u>T</u>ransporter isoform <u>4</u> (GLUT4) from the adipocyte vesicular pool to the cell surface is tightly regulated by insulin to maintain glucose homeostasis in response to changing demands in energy consumption. Previously, endosomal Na^+^/H^+^ exchanger 6 (NHE6, gene name *SLC9A6*) was identified in adipocyte plasma membranes and shown to associate with GLUT4-positive vesicles. However, the functional contribution of NHE6 to GLUT4 trafficking and glucose uptake remained uncharacterized.

**Methods:** NHE6 was knocked down in 3T3L1 cells either before or after differentiation into adipocytes using lentiviral delivery of shRNA. Uptake of [3H] 2-deoxyglucose into adipocytes was quantified in response to insulin. We used epitope tagged constructs to distinguish between localization of vesicular and surface levels of NHE6 and GLUT4 proteins. Protein and transcript levels of components of the glucose signaling pathway were monitored by qPCR and Western analysis, respectively, in response to NHE6 knockdown and/or treatment with chemical modulators of endosomal pH.

**Results:** We show that insulin-stimulated glucose uptake in adipocytes is severely impaired upon NHE6 depletion. Correspondingly, insulin-stimulated surface expression of GLUT4 at the adipocyte plasma membrane was diminished in NHE6 knockdown cells due to a post-transcriptional decrease in basal GLUT4. Metformin response of GLUT4 was also muted in the absence of NHE6. Further, we demonstrate diminished activation of the GLUT4 translocation pathway in the absence of NHE6 via reduced expression of the insulin receptor and reduced phosphorylation of the downstream effector kinase Akt. Components of GLUT4 storage vesicles, including GLUT4, LRP1 and sortilin were downregulated in NHE6-knockdown adipocytes under basal conditions. Proteostatic control of key components of the insulin signaling pathway (insulin receptor, GLUT4) could be restored by chemical bypass of NHE6 using the V-ATPase inhibitor bafilomycin or the Na^+^/H^+^ exchanger mimetic monensin.

**Conclusions:** Thus, NHE6 is critical for proper expression and trafficking of GLUT4. Basal expression of both insulin receptor and GLUT4 in NHE6 knockdown cells could be restored by bafilomycin-inhibition of the H^+^-ATPase or the H^+^ ionophore monensin pointing to pH dysregulation as the underlying defect. We suggest that NHE6 is a component of GLUT4 storage vesicles where it regulates proteostasis. These findings establish NHE6 as a novel contributor to glucose homeostasis and energy metabolism with implications for Christianson syndrome patients who carry loss of function mutations in the *SLC9A6* gene.

**Graphical Abstract:** 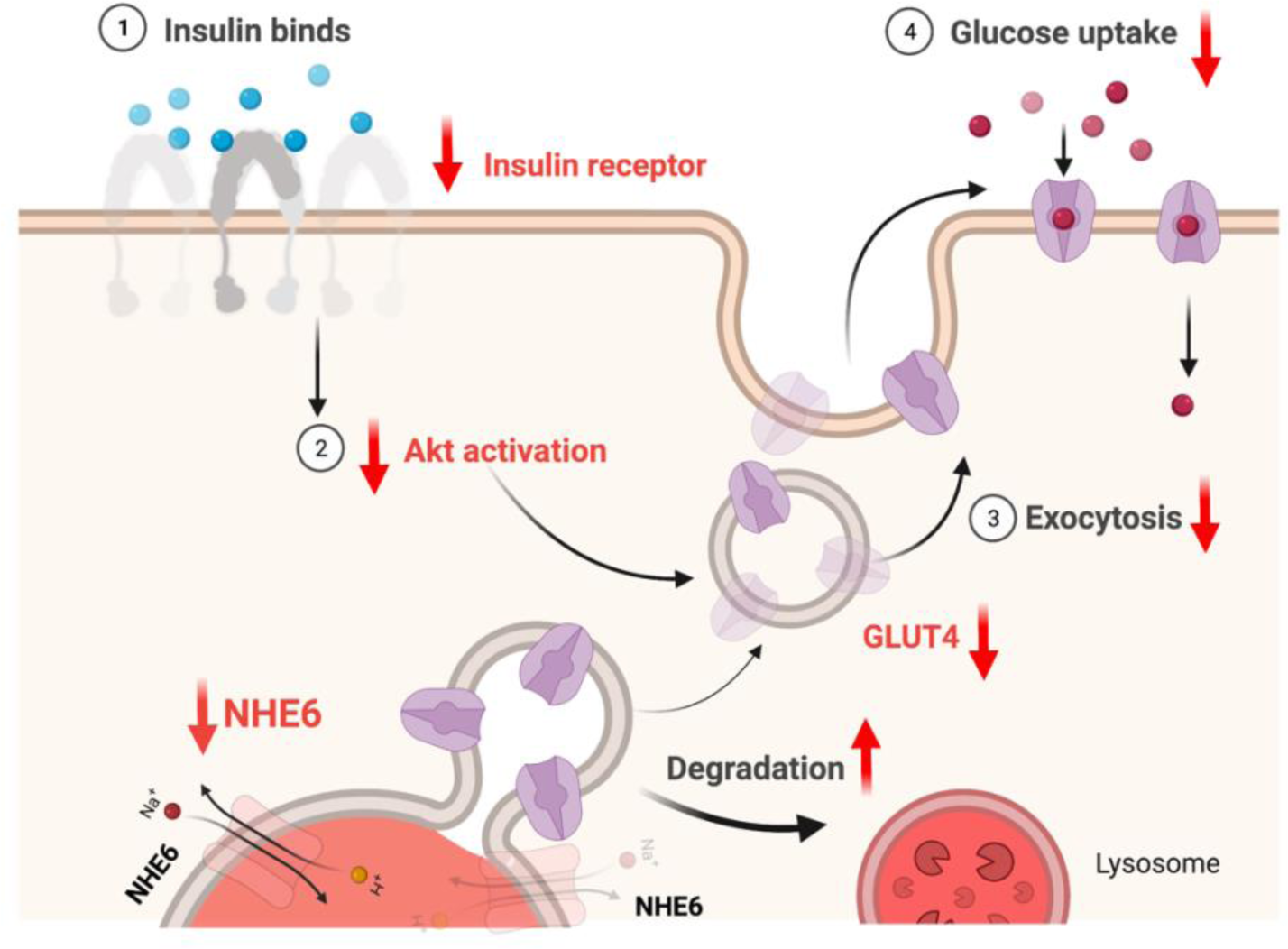

## 1. Introduction

Glucose homeostasis is maintained by insulin-responsive glucose uptake into muscle and fat. Insulin stimulates the rapid translocation of <u>GLU</u>cose <u>T</u>ransporter isoform <u>4</u> (GLUT4) from intracellular GLUT4 storage vesicles (GSV) to the plasma membrane [1]. Failure in insulin control of glucose transport is a chief component underlying insulin resistance Type 2 diabetes [2, 3]. GLUT4 expression is reduced in obese, prediabetic, insulin resistant, and diabetic individuals. Biguanide drugs such as metformin are widely used to treat diabetes and work, in part, by increasing glucose uptake into adipose tissue [4]. Specifically, metformin increases insulin-independent plasma membrane expression of GLUT4 in adipocytes under basal conditions [5]. Thus, many lines of independent evidence point to glucose uptake by GLUT4 as the rate limiting step in glucose homeostasis. Identifying the factors that influence GLUT4 expression and trafficking could lead to a better understanding of insulin resistance in diabetes which is essential for more effective management of this disease.

Endosomal pH plays a key role in regulating the trafficking and turnover of plasma membrane receptors and transporters. In adipocytes, alkalization of the endosome by the H^+^-ATPase inhibitor bafilomycin was shown to mimic insulin action by eliciting an immediate translocation of GLUT4 to the plasma membrane [6]. There are two ubiquitously expressed Na^+^/H^+^ exchanger isoforms, NHE6 and NHE9, that alkalinize the lumen of early and recycling endosomes by replacing luminal protons with cations [7]. Previous work has shown that gain or loss of endosomal NHE (eNHE) activity results in corresponding up or down regulation of plasma membrane receptors and transporters with significant and wide-ranging impact on tumorigenicity [8, 9], neurological function [10, 11], and cyst formation [12], to name a few examples. However, their role in GLUT4 expression and translocation has not been examined.

Evidence supporting a critical role for mammalian eNHE in glucose uptake has been observed across model organisms. Loss of eNHE orthologs in *C. elegans* and *Drosophila* confers metformin resistance, measured by survival and resistance to metformin toxicity [13, 14]. Consistent with the possibility that eNHE activity is required for metformin’s action, the ability of metformin to block starvation-induced autophagy is lost in eNHE knockout worms [13]. Starvation-induced upregulation of NHE6 is uniformly observed in many organisms and mammalian cells where it may be important to regulate autophagy, lysosome biogenesis or vesicle trafficking to promote survival [15]. A proteomic analysis of the adipocyte plasma membrane revealed NHE6 as a novel insulin-responsive protein [16]. Taken together, these observations suggest that the regulation of endosomal pH by eNHE may be important for insulin-stimulated glucose uptake yet the mechanistic basis for these intriguing observations is unclear.

In this study, we investigate a role for NHE6 in insulin-stimulated GLUT4 translocation in 3T3-L1 derived adipocytes. We show that NHE6 is required for insulin-stimulated glucose uptake by exercising proteostatic control on basal levels of several components of GLUT4 storage vesicles and downstream activators of the insulin-signaling pathway. Proteostatic control could be restored with chemical agents that alter the endosomal lumen, pointing to a role for endosomal pH regulation in glucose uptake.

## 2. Materials and Methods

### 2.1 Cell lines and culture

3T3-L1 fibroblasts (gifted from the lab of Dr. G. William Wong, The Johns Hopkins University School of Medicine), were maintained in DMEM (ThermoFisher Scientific, Waltham, MA) containing 10% Bovine Calf Serum (Sigma Aldrich, St. Louis, MO). Cells were maintained at 30% confluence at all times. Fibroblasts were differentiated into adipocytes using methods adapted from a protocol previously described[17]. 3T3-L1 fibroblasts containing stable expression of myc-GLUT4-mCherry [18, 19] were maintained and differentiated as above. Mature 3T3-L1 adipocytes were maintained in DMEM containing 10% Fetal Bovine Serum (Sigma Aldrich) until used for experimentation.

### 2.2 RNA extraction and qRT-PCR

Total RNA was extracted using QIAGEN’s RNeasy mini kit (QIAGEN, Germantown, MD) according to the manufacturer’s instructions. 1 μg of RNA was converted to cDNA using the High-Capacity RNA to cDNA kit (Applied Biosystems, Foster City, CA). For qRT-PCR, 50 ng of cDNA was added to master mix (Thermo Fisher Scientific), and probes *Slc9a6* (Mm00555445_m1), *Slc2a4* (Mm00436615_m1), *Adipoq* (Mm00456425_m1), *Plin1* (Mm00558672_m1), *Plin2* (Mm004757_m1), *Slc9a9* (Mm00626012_m1), *Gapdh* (Mm99999915_g1), *Insr* (Mm01211875_m1), *Slc2a1* (Mm00441480_m1) as specified.

### 2.3 Lentiviral transduction

NHE6-GFP in FuGW and shRNA constructs were transfected using lipofectamine 3000 (Invitrogen, Carlsbad, CA) into HEK293-FT cells to be packaged into virus using pCMV-Δ8.9 and PMDG at a ratio of 9:8:1. NHE6 shRNA sequence (5’-CCGGCGTCCTAGTGCATGTCTCTGTTCAAGAGACAGAGACATGCACTAGGACTTTTTC-3’) was designed to target 3’ untranslated region (3’UTR). The non-targeting shRNA sequence used as negative control was 5’-CAACAAGATGAAGAGCACCAA-3’ (Sigma-Aldrich). Lentivirus was collected beginning 48-hrs post-transfection and concentrated with Lenti-X Concentrator (Takara Bio USA, Mountain View, CA) for 3 consecutive days. Cells were transduced with lentivirus in Opti-MEM Reduced Serum medium (Thermo Fisher Scientific) for 48 hrs. For shRNA knockdown, cells were selected with puromycin at 10 μg/ml, based on kill curve data for cell line. Knockdown was confirmed by qRT-PCR.

### 2.4 Immunofluorescence

#### 2.4.1 Cell surface labeling

Mature adipocytes (differentiation day 8) were serum starved for 2 hrs at 37^°^C, 5% CO_2_. Cells were rinsed once in ice cold 1X DPBS (Quality Biological, Gaithersburg, MD) and fixed in ice cold 2% paraformaldehyde for 5 minutes at room temperature. Following a 5 min wash in PBS with 10 mM glycine, primary antibodies: anti-c-Myc (Invitrogen, MA1-980), anti-HA (Cell Signaling Technology, #3724), were added to unpermeabilized cells at room temperature for 15 minutes. After subsequent washes in PBS 10 mM glycine, cells were incubated with secondary antibodies anti-mouse Alexa Fluor 488 (Invitrogen, A11001), anti-mouse Alexa Fluor 555 (Invitrogen, A32727) for 20 minutes at room temperature. Following washes in PBS with 10 mM glycine, glass coverslips were mounted to glass slides using DAKO mounting medium (Agilent, Santa Clara, CA) and stored at 4^°^C until analysis.

#### 2.4.2 Internal labeling

Mature adipocytes (differentiation day 8) were serum starved for 2 hrs. Cells were rinsed once in ice cold 1X DPBS (Quality Biological) and fixed in ice cold 2% paraformaldehyde for 5 minutes at room temperature. Following a 5 min wash in PBS with 10 mM glycine, cells were permeabilized using 0.1% saponin in PBS + 10 mM glycine at room temperature for 10 minutes. Primary antibody anti-mCherry (Cell Signaling Technology, E5D8F) was added at room temperature for 15 minutes with washes in PBS 10 mM glycine to follow. Cells were incubated with secondary antibody anti-rabbit Alexa Fluor 633 (Invitrogen, A21070) for 20 minutes at room temperature. Following washes in PBS 10 mM glycine, glass coverslips were mounted to glass slides using DAKO mounting medium (Agilent) and stored at 4^°^C until analysis.

### 2.5 Microscopy and image quantification

Images were acquired using Zeiss LSM 700 laser scanning confocal microscope with 63X/1.4 Oil DIC M27X objective and illumination lasers at 405, 488, 561, and 633 nm. CZI images were loaded into *ZEN 2.3* (blue edition) to process 2-D data. To quantify surface localization, CZI images were loaded into *Imaris 9.5.1*. Plasma membrane renderings using the Surfaces function on 3-D data were created followed by mean intensity value calculations. For 3D reconstructions, the Spots function was used to represent voxels at the cell surface. For co-localization analysis, the Pearson’s correlation coefficient was used to quantify overlap.

### 2.6 2-Deoxy-D-Glucose uptake assay

Mature adipocytes (differentiation day 8) were serum starved in DMEM for 2 hrs. Cells were incubated with 100 nM insulin in KRH buffer (140 mM NaCl, 5 mM KCl, 1.3 mM CaCl_2_, 1.2 mM KH_2_PO_4_, 2.5 mM MgSO_4_, 5 mM NaHCO_3_, 25 mM Hepes, and 0.1% BSA) for 20 minutes. 2-deoxy-D-[1,2-^3^H] glucose (0.1 μCi) in KRH buffer containing 100 nM (or 0 nM insulin for no insulin controls) was added to each well for 10 minutes. After uptake, cells were washed 3 times in 1X PBS and lysed overnight in NaOH. Radioactivity was measured by liquid scintillation counting.

### 2.7 Protein extraction and Western blotting

Cells were collected and lysed in RIPA buffer (Thermo Fisher Scientific) supplemented with protease inhibitor cocktail (Cell Signaling), and phosphatase inhibitors (Sodium Fluoride and Sodium Orthovanadate, Sigma-Aldrich). After 30 minutes of incubation, cells were centrifuged at 13 000 × g for 30 minutes. Supernatant was collected and passed through 70m µM cell strainer to get rid of the debris. Protein quantification was done by the bicinchoninic acid assay. Twenty µg of protein in each sample was resolved by electrophoresis using 4-12% Bis-Tris gels (Thermo Fisher Scientific) and transferred to nitrocellulose membrane (Bio-Rad, Hercules, CA). Immunoblots were probed with antibodies (1:1000) from Cell Signaling Technologies: Insulin receptor-β (23413S); Glut4 (2213S); P-Akt (#4060P); Akt (#9272S); β-actin (4967S); Sortilin (#20681S) and from Invitrogen: LRP-1 (#PA5-81212); GAPDH (#MA5-15738), followed by incubation with HRP-conjugated secondary antibodies. Proteins were visualized using chemiluminescence substrate. Blots were analyzed using ImageJ.

### 2.8 Rescue of compartmental pH

Mature control or NHE6KD adipocytes were incubated with either 1 µM monensin (BioLegend, San Diego, CA) for 16 hrs or 25 nM bafilomycin (InvivoGen, San Diego, CA) for 2 hrs and following incubation adipocytes were collected and total protein was extracted.

### 2.9 Oil Red O Staining

Mature adipocytes (differentiation day 8) were washed twice with 1X PBS and fixed with 4% paraformaldehyde for 1 hr at room temperature. Cells were incubated in 0.5% Oil Red O solution in 60% isopropanol (v/v) for 1 hr at room temperature. Following 2 washes in 1X PBS, cells remained in 1X PBS at 4°C until imaged. Images were captured using Keyence BZ-X710 All-in-One Fluorescence Microscope.

### 2.10 Statistical Analysis and Graphs

Statistical analyses were performed using GraphPad Prism 9.3.1. Unpaired Student two-tailed *t* tests were used to determine statistical significance of the data. Results were considered significant at *P* <0.05. Significance and non-significance were shown as ^ns^ *P* > 0.05, \**P*<0.05, \*\**P*<0.01, \*\*\**P*<0.001, \*\*\*\**P*<0.0001. Biorender was used for drawing schematics and panels were assembled in Adobe Illustrator.

## 3. Results

### 3.1 Differentiated adipocytes selectively upregulate NHE6

3T3-L1 fibroblasts were differentiated using an adipogenic cocktail adapted from methods described previously [17], and successful differentiation was confirmed by Oil Red O staining of cells eight days after induction of differentiation (**Fig. 1A**). An increase in mRNA of canonical adipocyte genes *Slc2a4* (GLUT4), *Adipoq* (Adiponectin) and *Plin1* (Perilipin-1) served as further confirmation of successful differentiation. However, the increase in *Plin-2* (Perilipin-2) was not as large as previously described [20] (**Fig. 1B**). A nearly 3-fold increase in NHE6 transcript was observed in 3T3-L1 adipocytes when compared to undifferentiated fibroblasts (**Fig. 1C**). While the magnitude of NHE6 increase is not on the same scale as the adipocyte markers (**Fig. 1B**), the high rate of ion exchange (∼1500 ion/s) associated with this class of transporter [21] means that small changes in expression can have large consequences in the limited microenvironment of the endosomal compartment. In contrast, transcript levels of the closely-related NHE9 isoform decreased by about 50% upon adipocyte differentiation (**Fig. 1D**) suggesting an isoform-selective role for NHE6 in adipocyte function.

**Figure 1.**
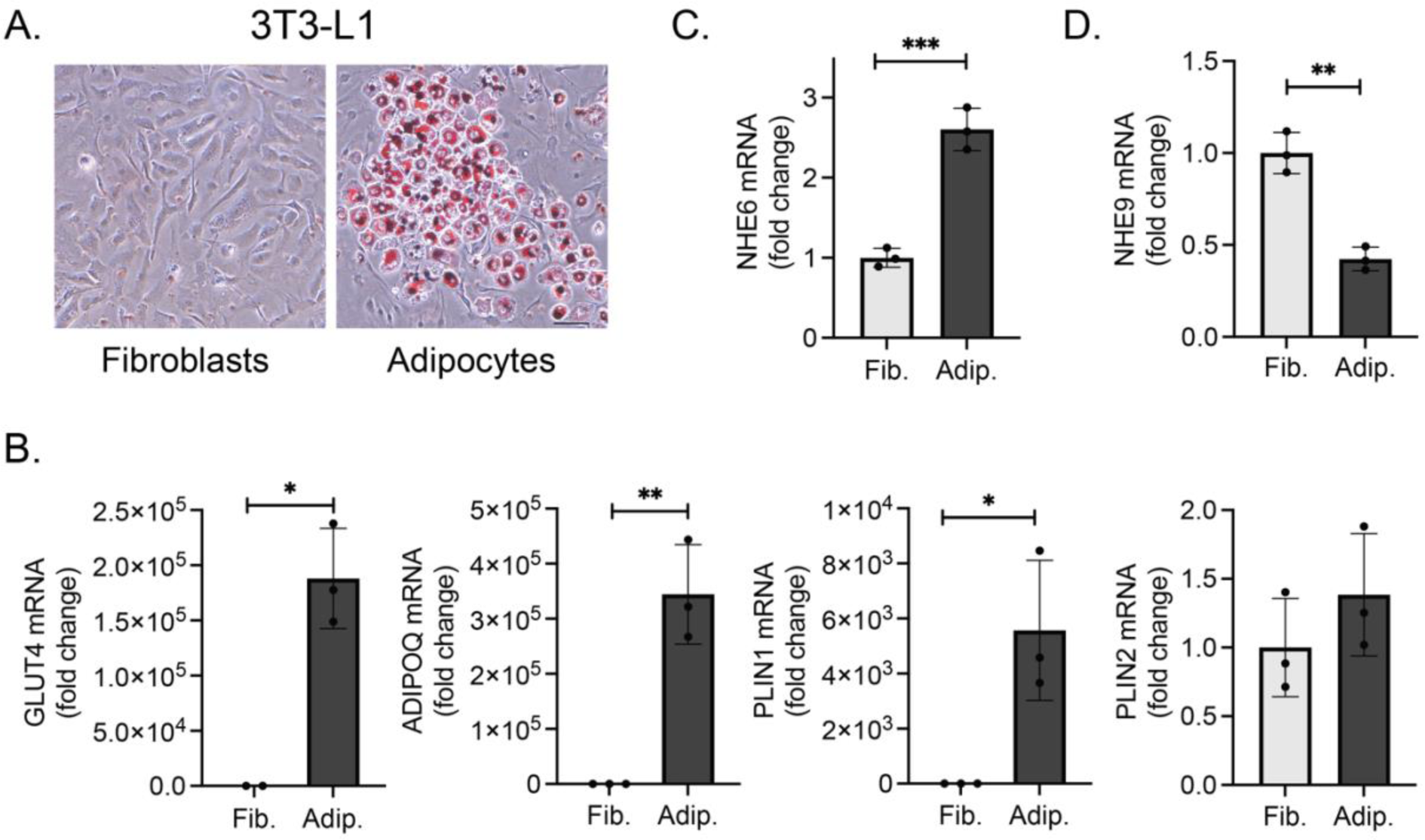
Differentiated 3T3-L1 adipocytes selectively upregulate NHE6. **(A)** 3T3-L1 fibroblasts (left) were differentiated into adipocytes (right) and stained with Oil Red O. Images were taken at 10x on day 8 of differentiation (scale bar, 100 µm). **(B)** qPCR analysis of gene transcripts from 3T3-L1 fibroblasts and adipocytes, shown as fold change. Canonical adipocyte genes *Slc2a4* (GLUT4), *Adipoq* (Adiponectin), *Plin1* (Perilipin-1), and *Plin2* (Perilipin-2) increased post differentiation. **(C-D)** Transcripts of endosomal Na^+^/H^+^ exchanger isoforms were determined by qPCR. NHE6 expression **(C)** increased 3-fold upon differentiation whereas NHE9 transcripts **(D)** decreased more than 2-fold. Graph show mean ± SD from three independent experiments. Data were analyzed using *t-*test. *\*P<0.05, **P<0.01, ***P<0.001*.

### 3.2 Insulin treatment increases NHE6 on the plasma membrane

Insulin elicits dynamic molecular responses in adipocyte plasma membranes that are key for rapid uptake and clearance of glucose from blood [16]. To discern the functional significance of NHE6 upregulation in adipocytes, we compared the cellular localization of NHE6 under basal conditions and following insulin treatment. 3T3-L1 cells with stable expression of NHE6 containing an external triple-HA tag and cytoplasmic C-terminal GFP marker (**Fig. 2A**) were antibody labeled without permeabilization. This approach allows the plasma membrane-localized NHE6 (HA signal) to be distinguished from total NHE6 (GFP signal). Previous work has shown that NHE6 is predominantly localized to early and recycling endosomes [22], although plasma membrane expression has been reported in some cells [23, 24]. Consistent with these findings, localization of NHE6-GFP in 3T3-L1 adipocytes was intracellular under basal conditions (**Fig. 2B**). In the presence of insulin there was a 2.5-fold increase in cell surface NHE6 levels in 3T3-L1 adipocytes as revealed by labeling of the extracellular HA epitope (**Fig. 2B-C**). 3D visualization of confocal microscopy images further illustrates the shift in the cellular localization of the 3×-HA-NHE6-GFP pool from intracellular to surface membrane-associated in response to insulin treatment (**Fig. 2D**). The enrichment of NHE6 at the cell surface of insulin-treated adipocytes we observe is consistent with previous findings [16], supporting an emerging link between insulin signaling and NHE6 function.

**Figure 2.**
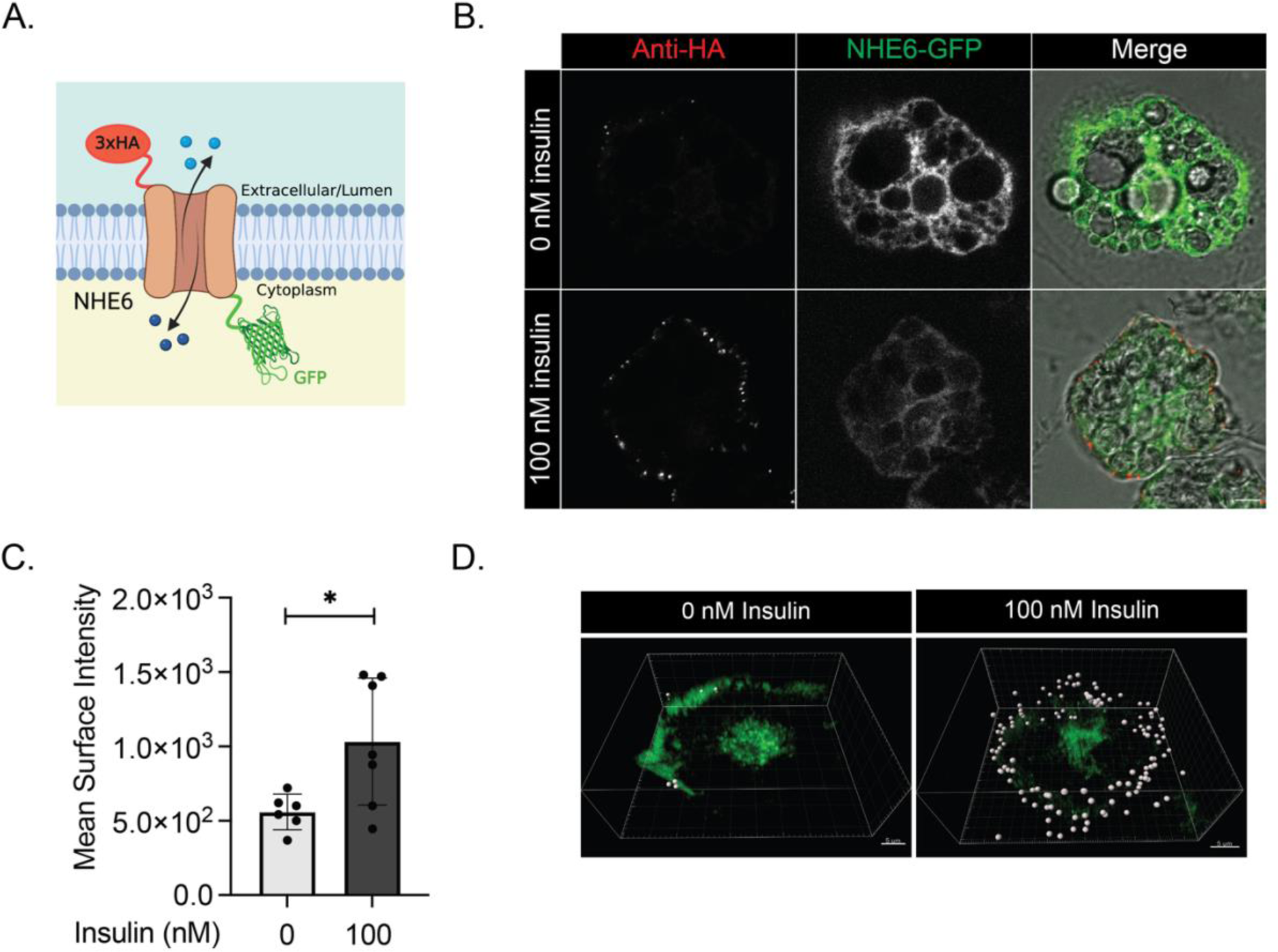
NHE6 is enriched at the plasma membrane in response to insulin. **(A)** Schematic of NHE6 construct extracellularly tagged with 3×HA and with GFP at the cytoplasmic C-terminus. **(B)** Representative immunofluorescence confocal images show increase of NHE6 at the cell surface of 3T3-L1 adipocytes following 100 nM insulin treatment when compared to untreated (0 nM) control cells. Confocal images show a single plane through the midpoint of each cell. **(C)** Quantification of surface expression of NHE6 assessed by anti-HA staining using Imaris. Graph shows mean surface intensity of NHE6 ± SD for n= 6 control and n= 7 insulin treated cells. Data were analyzed using *t*-test, *\*P<0.05*. **(D)** 3D visualization of control (0 nM) and insulin treated (100 nM) 3T3-L1 adipocytes with labeling of the surface expression using the spots function in Imaris. All Images were taken at 63× magnification (scale bar, 5 µm).

### 3.3 NHE6 and GLUT4 have overlapping cellular localization

A consequence of insulin signaling is the global acceleration of exocytosis which can lead to an insulin-dependent increase of endosomal proteins at the cell surface [25–29]. We wanted to rule out the possibility that the surface enrichment of NHE6 reflects a generic increase in endosomal exocytosis and to determine whether there is a potential link between insulin-stimulated GLUT4 translocation and NHE6. Therefore, we investigated co-localization of NHE6 and GLUT4 under basal and insulin-stimulated conditions by expressing HA-NHE6-GFP in 3T3-L1 adipocytes stably transfected with a cMyc-GLUT4-mCherry construct (**Fig. 3A**) [18, 19]. In the absence of insulin, the majority of GLUT4 is distributed between endosomes, trans Golgi network and heterogeneous mix of endosomal sorting intermediates and specialized GLUT4 storage vesicles (GSVs) [30]. We observed moderate co-localization between NHE6-GFP and GLUT4-mCherry signals in the basal state (Pearson’s correlation, 0.32) (**Fig. 3B-C**). Following insulin treatment, the extent of co-localization remained the same (**Fig. 3B, D**). This finding is consistent with translocation of the co-localized intracellular pool of NHE6-GFP and GLUT4-mCherry to the plasma membrane.

**Figure 3.**
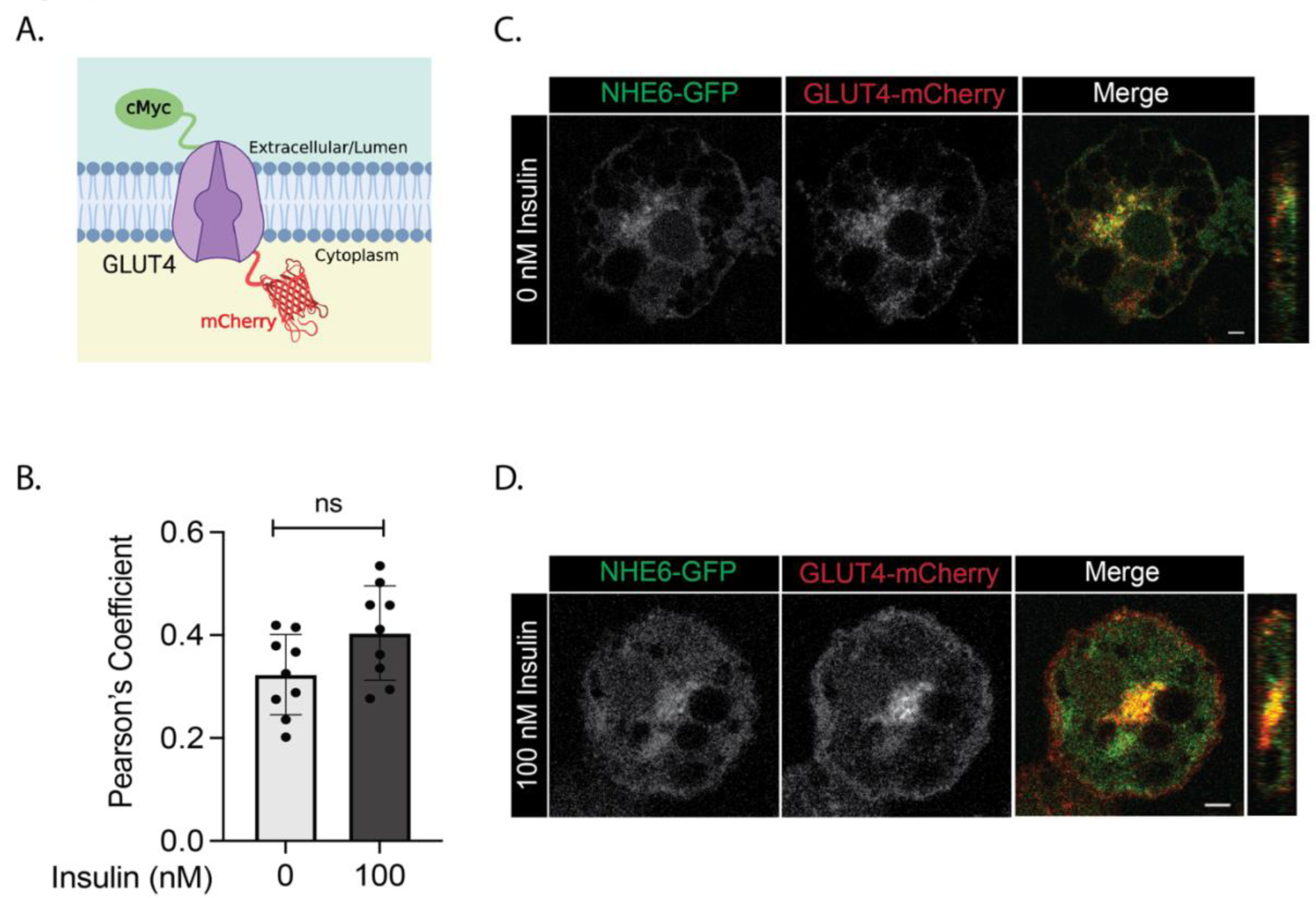
NHE6 and GLUT4 partially co-localize in intracellular vesicles. **(A)** Schematic of GLUT4 construct extracellularly tagged with cMyc epitope and mCherry at the cytoplasmic C-terminus. cMyc-GLUT4-mCherry was stably expressed in 3T3-L1 cells. (**B-D**) Co-localization of HA-NHE6-GFP (GFP signal) and cMyc-GLUT4-mCherry (mCherry signal) under basal conditions (serum starvation for 2 hrs) or after 100 nM insulin treatment in 3T3-L1 adipocytes. Confocal images show a single plane through the midpoint of each cell. Images were taken at 63× magnification (scale bar, 5 µm). Pearson’s correlation coefficient computed from n=9 cells. Graph show mean ± SD; ns = not significant.

### 3.4 NHE6 knockdown decreases insulin-stimulated glucose uptake by reducing cell surface GLUT4

To evaluate the functional role of NHE6 in insulin-dependent GLUT4 translocation and glucose uptake, we used shRNA to knockdown (KD) NHE6 in 3T3-L1 cells either before or after differentiation into adipocytes. Although NHE6 knockdown in 3T3-L1 fibroblasts led to a reduction in the total number of adipocytes post-differentiation (**Fig. S1A**), expression of canonical adipocyte genes was comparable to control (**Fig. S1B**). We confirmed that transcript levels of the related endosomal NHE isoform, NHE9, remained unchanged upon NHE6 knockdown (**Fig. S1C**).

Insulin-responsive glucose uptake was monitored in control and NHE6 knockdown 3T3-L1 adipocytes (**Fig. 4A**). In control adipocytes, glucose uptake was increased by 7-fold following acute treatment with insulin as previously reported [1] (**Fig. 4B**). In contrast, NHE6 KD adipocytes showed only a 2-fold increase in glucose uptake in response to insulin (**Fig. 4B**).

**Figure 4.**
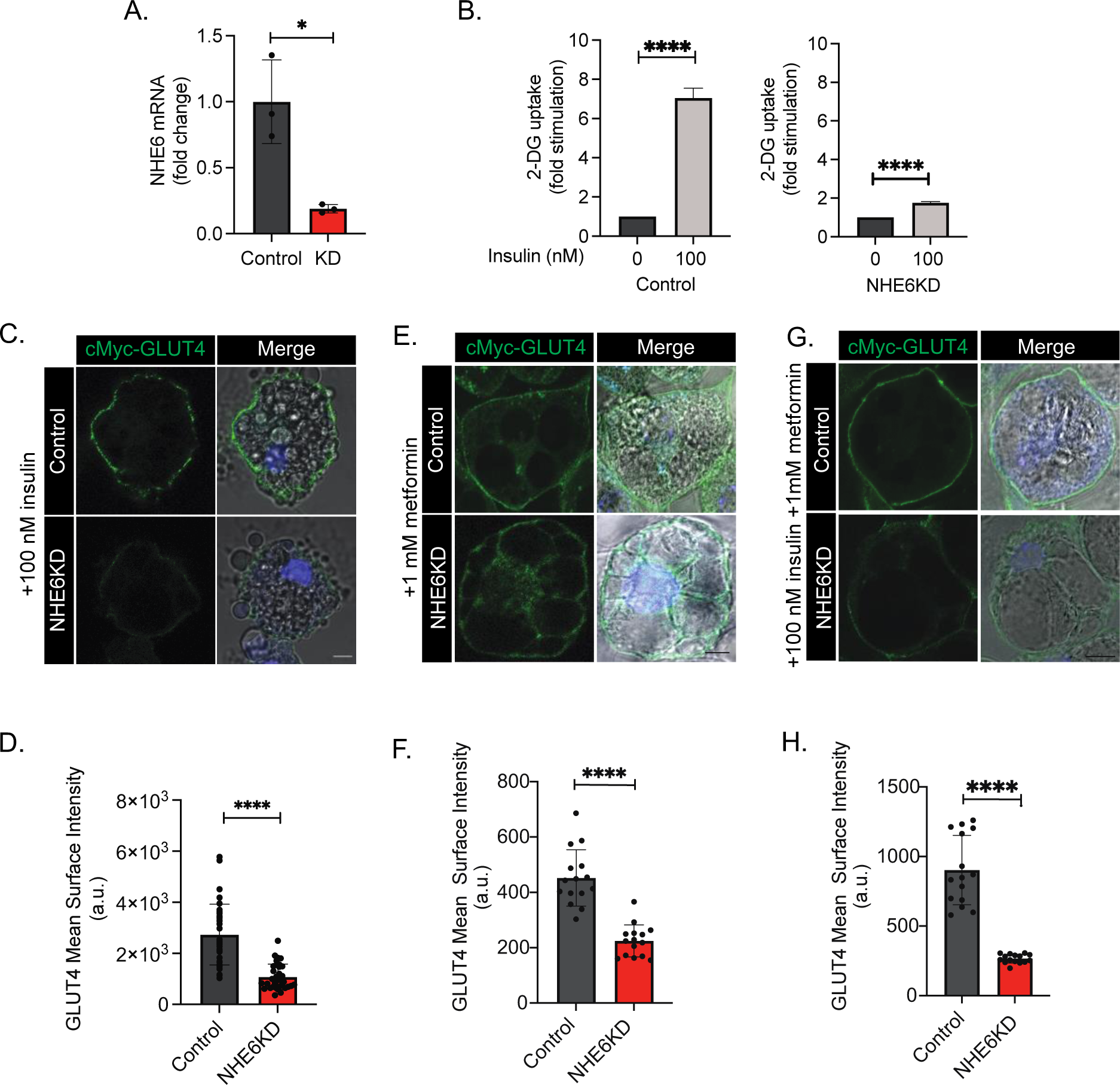
NHE6 is required for insulin-stimulated glucose uptake and surface expression of GLUT4 in adipocytes. **(A)** NHE6 knock down in 3T3-L1 adipocytes. **(B)** Insulin-stimulated glucose uptake is impaired by NHE6 knockdown. Differentiated adipocytes were serum starved, treated with 100 nM insulin, and assayed for 2-deoxyglucose uptake (CPM lysate/ µg protein) for 5 min. Uptake was normalized to respective 0 nM insulin controls. Graphs show mean ± SD for n=6 biological replicates per condition. **(C)** Representative images and **(D)** quantification of the mean intensity of cMyc-GLUT4-mCherry surface labeling (Myc signal) showing reduction of GLUT4 at the cell surface of NHE6 KD 3T3-L1 adipocytes following 100 nM insulin treatment. **(E)** and **(F)** Cells were treated for 24 h with 1 mM metformin or **(G)** and **(H)** with 1 mM metformin for 24 h followed by 100 nM insulin. All confocal images show a single plane through the midpoint of each cell. Images were taken at 63× magnification, scale bar 5 µm. Quantification was done by Imaris **(D)** or Image J **(F, H)**. Graph shows mean ± SD for n= 15-34 cells per group. Data were analyzed using *t-*test*. *P<0.05,* \*\*\**\*P<0.0001*.

NHE6 regulates cell surface trafficking of transmembrane receptors and transporters in other cell types [10, 11]. Thus, we next determined whether changes in glucose uptake in NHE6 KD adipocytes were caused by defects in trafficking of GLUT4 in response to insulin. We treated control and NHE6 KD adipocytes expressing c-Myc-GLUT4-mCherry with 100 nM insulin and labeled the surface GLUT4 pool using an anti-c-Myc antibody in non-permeabilized cells (**Fig. 4C**). NHE6 knockdown resulted in a nearly 66% reduction in the amount of GLUT4 at the adipocyte cell membrane (**Fig. 4D**). Insulin-dependent GLUT4 cell surface localization was similarly diminished in mature NHE6 KD adipocytes in which gene knockdown was induced post-differentiation (**Fig. S1D-E**). The robust downregulation of GLUT4 surface expression in insulin-treated NHE6 KD adipocytes likely underlies the observed decreases in insulin-mediated glucose uptake. We also evaluated insulin-independent surface expression of GLUT4 by treating adipocytes with 1 mM metformin for 24 h. NHE6 knockdown also diminished metformin-induced GLUT4 surface labeling, both in the absence (**Fig. 4 E-F**) or presence (**Fig. 4 G-H**) of additional treatment with insulin (30 min, 100 nM).

### 3.5 NHE6 regulates proteostasis and activation of the GLUT4 pathway

To better understand how NHE6 regulates GLUT4 membrane transport and glucose uptake in adipocytes, we evaluated protein levels of GLUT4 and other components of the insulin signaling pathway in adipocytes. Surprisingly, mature NHE6 KD 3T3-L1 adipocytes express 44% less GLUT4 (**Fig. 5A, B**) under basal conditions despite no significant changes in GLUT4 transcript levels (**Fig. S2A**). These data indicate that decreases in insulin-stimulated GLUT4 accumulation at the cell surface of NHE6 KD adipocytes are due, in part, to a post-transcriptional reduction in basal levels of GLUT4 protein. In contrast, protein levels of GLUT1, the transporter responsible for the bulk of basal glucose uptake, increased by ∼60% in NHE6 KD adipocytes, also without changes in mRNA expression (**Fig. S2B-D**). This observation points to a selective effect of NHE6 in the regulation of GLUT4 levels and glucose homeostasis in adipocytes.

**Figure 5.**
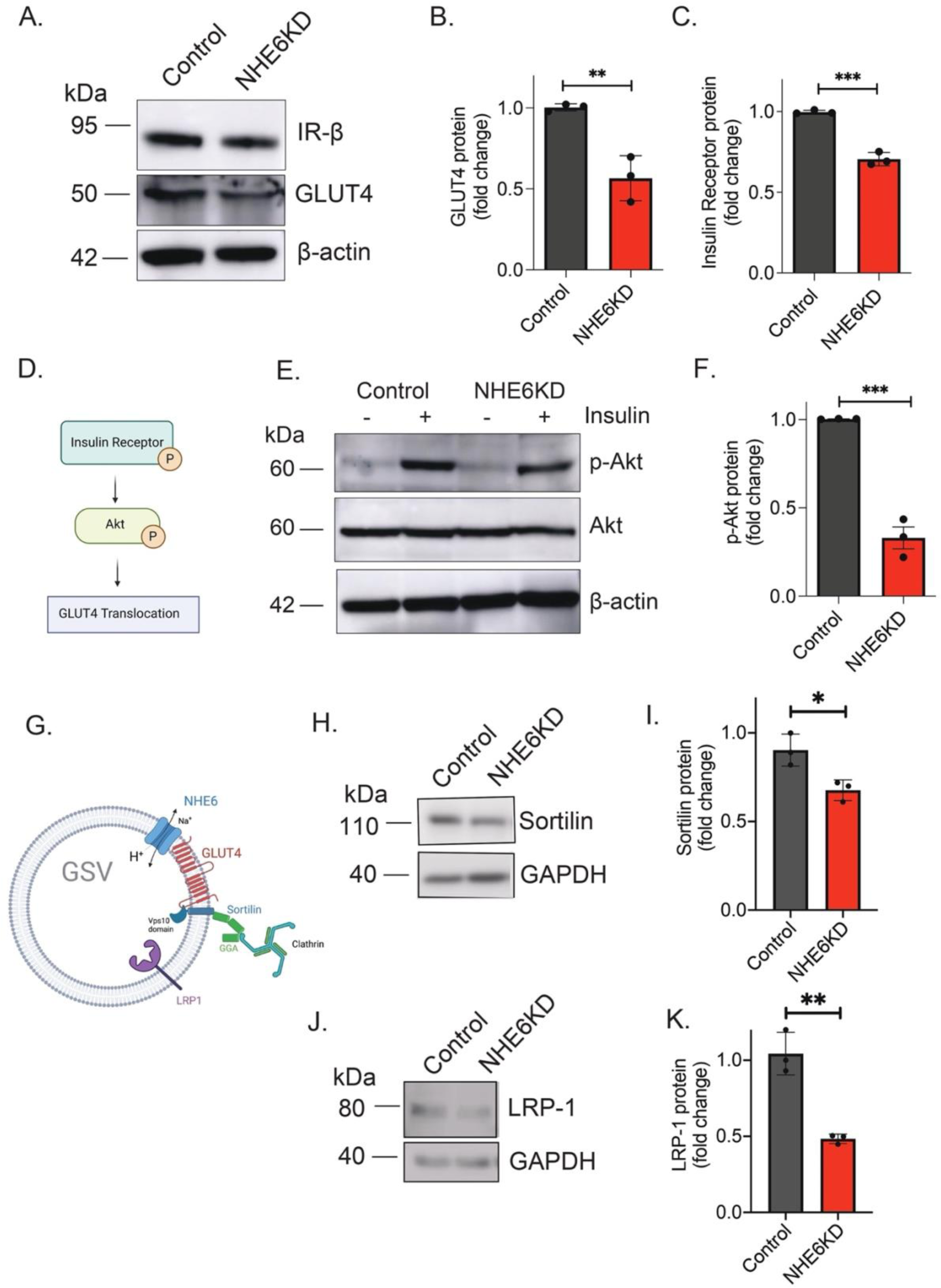
NHE6 regulates proteostasis and activation of the GLUT4 pathway. **(A)** Representative western blot shows total levels of insulin receptor (IR-β) and GLUT4 proteins in control and NHE6 KD 3T3-L1 adipocytes under basal conditions. β-actin was used to as a loading control. **(B)** Quantification of normalized total protein levels. NHE6 reduction significantly decreased GLUT4 (-0.44 ± 0.08) levels in comparison to control adipocytes. **(C)** Knockdown of NHE6 significantly decreased insulin receptor total protein levels (-0.29 ± 0.02) in comparison to control adipocytes. **(D)** Schematic of Akt activation by phosphorylation. **(E)** Representative western blot shows total levels of p-Akt ^Ser473^ and Akt in control cells and NHE6 KD adipocytes at the basal state and upon insulin treatment. NHE6 loss reduces pAkt ^Ser473^ (-0.67 ± 0.06) but not total Akt levels. **(F)** NHE6 KD adipocytes have significantly reduced p-Akt^Ser473^ levels when compared to control adipocytes. **(G)** Cartoon of GLUT4 Storage Vesicle (GSV) showing major protein components (GLUT4, sortilin, and LRP1) and proposed co-localization of NHE6. **(H, J)** Representative western blots show sortilin **(H)** and LRP1 **(J)** in adipocyte lysates and their quantitation **(I-K)**. NHE6 knockdown significantly decreased sortilin (-0.32 ± 0.05) and LRP1 levels (-0.52 ± 0.03) in comparison to control adipocytes. Graphs show mean ± SD for n=3 biological replicates/group. Data were evaluated using *t*-test. \**P<0.05,* \**\*P<0.01, ***P<0.001*

In line with their lessened glucose uptake in response to insulin, NHE6 KD adipocytes also showed a 30% reduction in protein levels of the insulin receptor-β (IR-β) subunit (**Fig. 5A, C**), even though transcript levels were increased by 24% (**Fig. S2E**). Therefore, we investigated whether reduced insulin receptor levels affect downstream activation of the insulin signal transduction pathway. The serine/threonine kinase Akt is a key downstream effector of insulin-stimulated GLUT4 translocation (**Fig. 5D**) [31]. Akt activation alone is sufficient to stimulate GLUT4 translocation in the absence of insulin [32]. We found that phosphorylated Akt (p-Akt^Ser473^) levels were reduced 67% in insulin-treated NHE6 KD adipocytes without change in total Akt protein levels (**Fig. 5E-F**). We conclude that activation of this branch of the insulin signaling pathway is impaired in the absence of NHE6.

Next, we asked if other membrane protein components of the GSV show altered proteostasis in response to NHE6 knockdown. One key component of GSVs is sortilin, a Vps10 domain containing protein that interacts with GLUT4 within the endosomal lumen and recruits components of the retromer and clathrin with its cytoplasmic tail [33] (**Fig. 5G**). A proteomic analysis of GSVs revealed LRP1 (low density lipoprotein receptor-related protein 1) to be another major, insulin-responsive component [34]. We found that NHE6 knockdown decreased protein levels of both LRP1 and sortilin (**Fig. 5H-K**), similar to that of GLUT4. Taken together, our data demonstrate that NHE6 regulates proteostasis of multiple components of the GLUT4 storage vesicles and insulin signaling pathway.

### 3.6 Chemical bypass of NHE6

The exchange of luminal protons for cations by transport activity of NHE6 counters the proton pumping activity of the V-ATPase to establish the pH balance of the endosome [35, 36]. Thus, loss of NHE6 results in hyper-acidification of endosomal lumen by 1-2 pH units [10, 36]. To determine whether the effects of NHE6 loss on the GLUT4 translocation pathway are linked to aberrant compartmental pH, we attempted chemical rescue of NHE6 KD adipocytes with agents known to alkalinize endosomal pH. The ionophore monensin is a chemical mimetic of Na^+^/H^+^ exchangers and has been shown to block lysosomal delivery of cargo and reroute proteins to the cell surface [37]. Previously, we showed that monensin treatment reversed hyperacidic pH of endosomes resulting from NHE6 KD [9, 11, 36]. Western blot analysis of NHE6 KD adipocytes treated with 1µM monensin for 16 hr was sufficient to restore GLUT4 and insulin receptor proteins to levels similar to control adipocytes (**Fig. 6A-C**). Using a different mechanism to rescue lumenal pH, by treating with the V-ATPase inhibitor bafilomycin, we also restore protein levels of both insulin receptor and GLUT4 in NHE6 KD adipocytes (**Fig. S3**).

**Figure 6.**
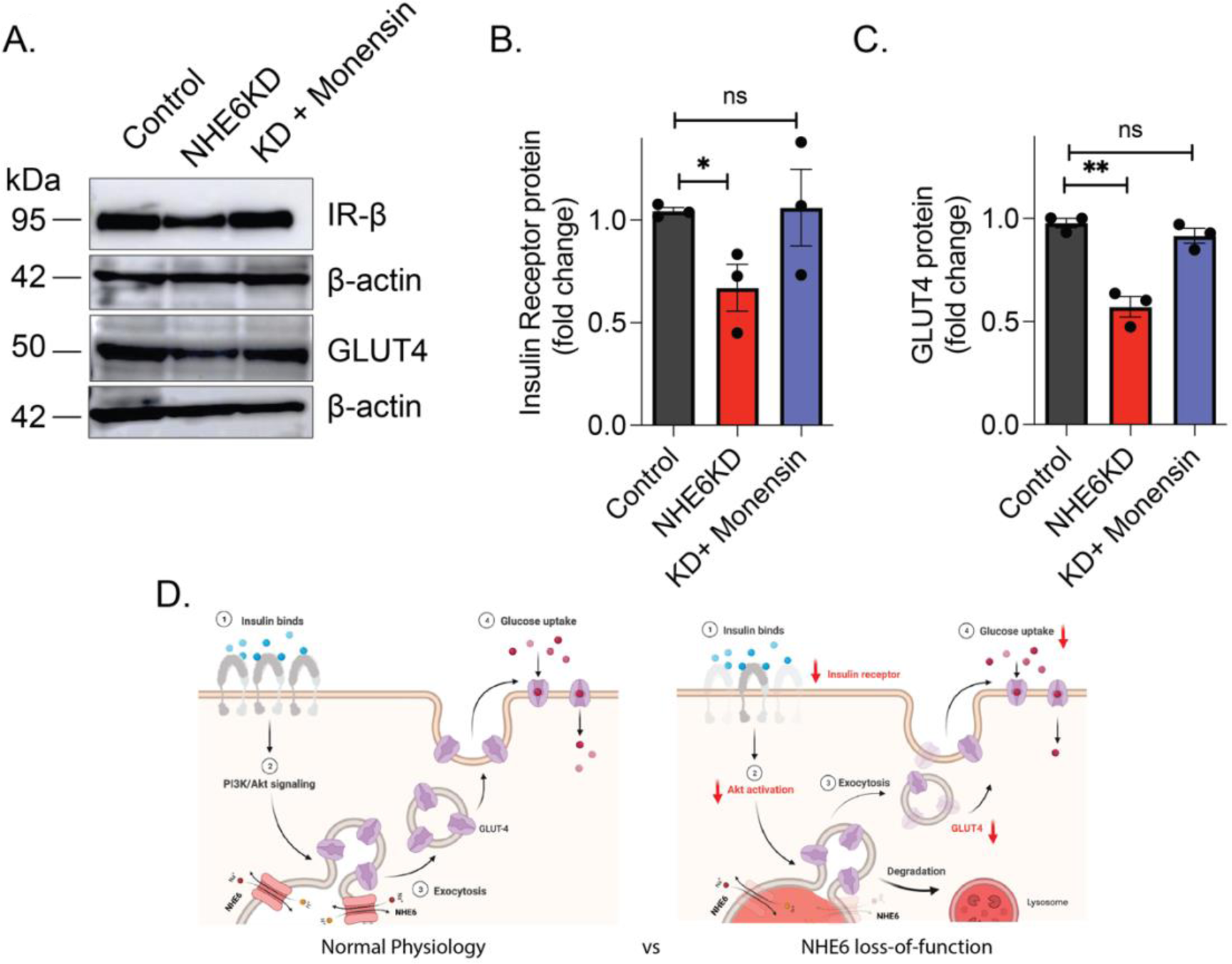
pH-modulating drugs rescue defects caused by NHE6 knockdown. **(A)** Representative western blot showing total levels of insulin receptor (IR-β) and GLUT4 in control, NHE6 KD, and NHE6 KD adipocytes treated with 1 µM monensin for 16 hrs. β**-** actin was used as a loading control.**(B)** Treatment with monensin, which increased luminal pH, restores IR-β protein expression to levels of control adipocytes. **(C)** Monensin treatment restores GLUT4 protein expression to levels of control adipocytes. Graphs in B and C show mean ± SD for n=3 biological replicates/group. Data were evaluated using *t-*test. *\*P<0.05,* \**\*P<0.01.* ns = not significant. **(D)** Schematic summarizing the effect of NHE6 loss on the GLUT4 translocation pathway.

In summary, these data point to multiple steps involved in adipocyte insulin-stimulated glucose uptake that are affected by loss of NHE6 (**Fig. 6D**). Taken together, pH regulation of the endo-lysosomal pathway is critical for maintaining GLUT4 levels in adipocytes and for its proper translocation to cell surface in response to insulin.

## 4. Discussion

In pre-adipocytes, both ectopically and endogenously expressed GLUT4 localizes to endosomes with a half-life of only 2 hours [33]. However, in differentiated adipocytes, GLUT4 is sorted into small (<100 nm) GLUT4 storage vesicles which represent the major insulin-responsive compartment. This not only increases its insulin responsiveness but also protects GLUT4 from lysosomal degradation. What controls this proteostatic switch? Evidence from this study reveals a critical role for pH inside the lumen of the endo-lysosomal compartments in controlling proteostasis of GLUT4 and other essential components of the GSV.

Previous work established NHE6 as a novel insulin responsive protein that associates with GLUT4-positive vesicles in adipocytes [16]. Our work confirms the enrichment of NHE6 upon adipocyte differentiation and at the adipocyte plasma membrane in response to insulin. However, the functional role of NHE6 in glucose homeostasis and GLUT4 trafficking had not previously been established. Our work shows that loss of NHE6 strongly reduces insulin-stimulated glucose uptake, due to a corresponding reduction of GLUT4 at the cell surface. Reduced surface expression of GLUT4 is caused in part by reduced activation of the insulin signaling pathway (evidenced by reduced expression of insulin receptor and Akt phosphorylation) and in part due to decreased basal levels of GLUT4 protein. Other studies report the existence of discrete organellar pools of GLUT4 within the cell with about 40% of the total residing in the endo-lysosomal system [1, 38]. Our study determined NHE6 loss corresponding to an approximately 44% decrease in total GLUT4 protein suggesting that NHE6 regulates the itinerary of GLUT4 through the endo-lysosomal system. These defects in GLUT4 stability and trafficking can be rescued when treating with monensin, a chemical mimetic for Na^+^/H^+^ exchange, suggesting the ion transport activity of NHE6 is required for this process.

Glucose uptake in adipose tissue accounts for 10% of whole-body glucose uptake [39]. Therefore, it is important to consider whether NHE6 regulates glucose availability in other cell-types and tissues. Given the effect NHE6 has on insulin receptor levels, the role of NHE6 in skeletal muscle function should also be explored. In the CNS, GLUT4 has been demonstrated to be enriched at active synapses likely supporting energy demands during sustained activity [40]. A question of interest is whether NHE6 also regulates glucose uptake at active synapses. Christianson syndrome is a severe neurological disorder resulting from mutations in NHE6 [41]. Low body weight is a common comorbidity observed in CS patients [41–43] and may be linked to NHE6 loss of function in adipose-derived stem cells or impairment of anabolic responses linked to insulin. *In vivo,* adipose-derived stem cells are highly sensitive to insulin and play an important role in tissue development and maintenance. We note that NHE6 knockdown in pre-adipocytes reduced the efficiency of differentiation into mature adipocytes and increased levels of adiponectin, possibly as a compensatory response to increase insulin sensitivity [44].

GLUT4 expression is reduced in obese, prediabetic, insulin resistant, and diabetic individuals [2, 39]. In mouse models, heterozygous knockout of GLUT4 and conditional deletion in adipose tissue or skeletal muscle results in insulin resistance and an inclination toward diabetes [45, 46]. GLUT4 gene variants are rare [47]; therefore, understanding its regulation is key to understanding metabolic disorders. The diabetic drug metformin increases the basal expression of GLUT4. Intriguing findings from worm and fly models show that eNHE mutants are resistant to metformin, likely through dysregulation of autophagy [13]. We show here that knockdown of NHE6 greatly reduced surface expression of GLUT4 in response to metformin. Thus, NHE6 may be a target of metformin and future studies should explore the role of NHE6 in GLUT4 regulation by metformin.

## 5. Conclusions

In summary, key events in the insulin signaling pathway in 3T3L1 adipocytes are regulated by the endosomal Na^+^/H^+^ exchanger, NHE6. We confirmed that NHE6 is upregulated in differentiated adipocytes where it partly co-localizes with GLUT4 in vesicular compartments and traffics to the adipocyte surface in response to insulin. We observed prominent reduction of Insulin Receptor levels and decreased insulin-responsive Akt phosphorylation following NHE6 knockdown. Several components of GLUT4 storage vesicles including GLUT4, LRP1 and sortilin were decreased in the absence of NHE6, resulting in decreased surface expression of GLUT4 in response to insulin. Together, these effects culminated in severe reduction in insulin-stimulated glucose uptake in NHE6 knock down adipocytes. GLUT4 response to metformin was also muted in the absence of NHE6, pointing to insulin-independent effects. Finally, we showed that inhibitors of lyso-endosomal acidification (bafilomycin) and chemical mimetics of Na^+^/H^+^ exchange (monensin) rescued protein expression of Insulin Receptor and GLUT4 in the absence of NHE6. Our findings reveal a critical role for NHE6 in pH-mediated proteostatic control of glucose uptake pathways in 3T3L1 adipocytes.

## Acknowledgements

We thank Reina Ambrocio for help with Biorender drawings. R.R. was supported by grants from the National Institutes of Health (R01GM147197) and BSF (13044). D.N.L. was supported by a grant from the American Diabetes Association (1-19-JDF-081). A.X.M. was a Gilliam Fellow of the Howard Hughes Medical Institutions and M.R.M was an AACR-AstraZeneca Breast Cancer Research Fellow. R.W.B. acknowledges support from Formation Venture Engineering. C.J.W. was supported by F99NS118713 and K00NS118713 from the National Institutes of Health. A.X.M. and C.J.W. were supported by T32GM144272 to the Biochemistry, Cellular and Molecular Biology Program at Johns Hopkins University School of Medicine from the National Institutes of Health.

---

**Figure S1.**
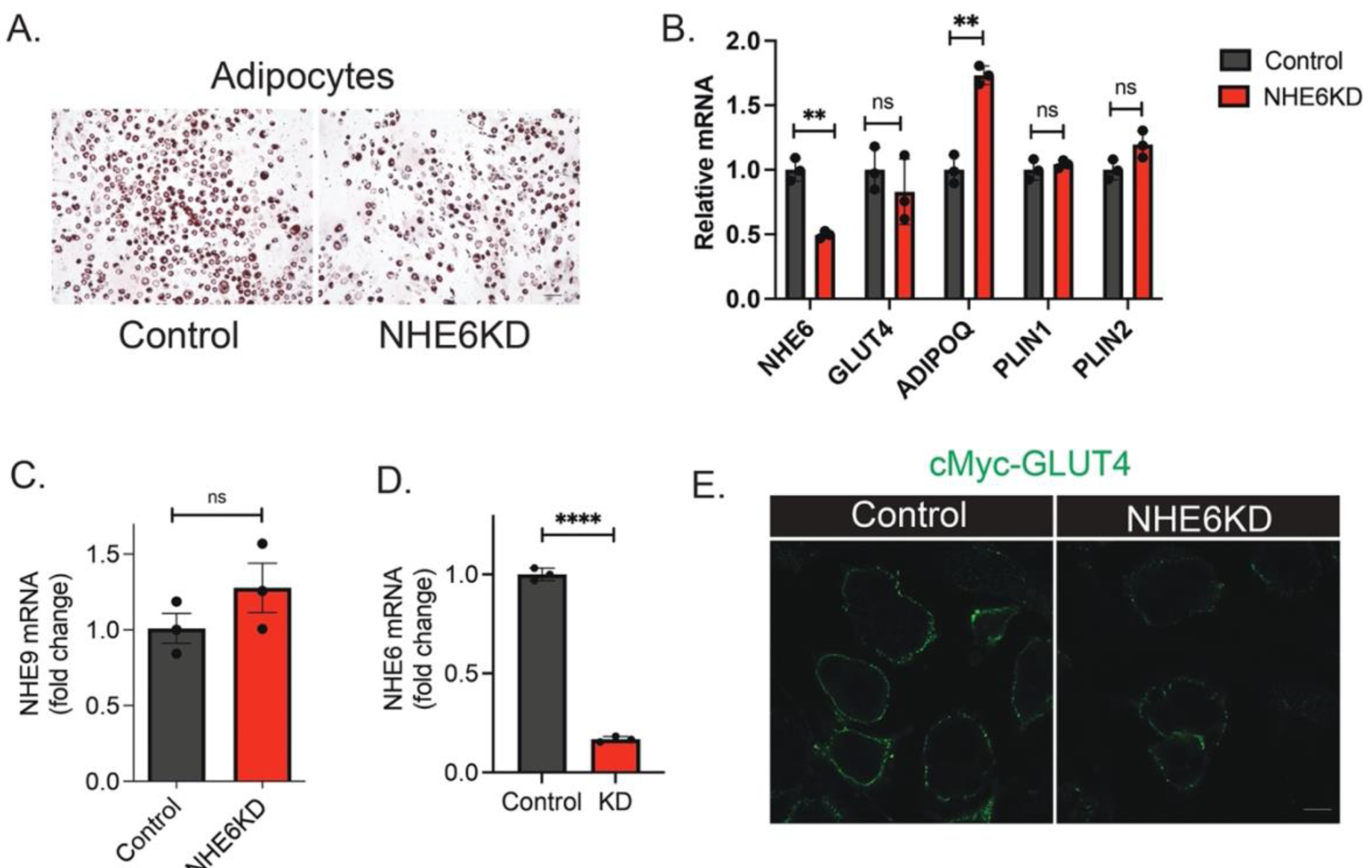
Comparing knockdown of NHE6 pre and post differentiation. **(A)** Oil Red O staining of control and NHE6 KD 3T3-L1 adipocytes eight days post-adipocyte differentiation. Cells were treated with shRNA pre-differentiation. Knockdown of NHE6 pre-differentiation causes a qualitative reduction in the density of 3T3-L1 adipocytes. Images were taken at 10× magnification, (scale bar 100 µm). **(B)** mRNA expression of adipocyte genes *Slc2a4* (GLUT4), *Adipoq* (Adiponectin), *Plin1* (Perilipin-1), and *Plin2* (Perilipin-2) in 3T3-L1 adipocytes treated with shRNA pre-differentiation (scramble or shNHE6). NHE6 KD adipocytes show similar gene expression to control adipocytes, except for increased adiponectin levels. **(C)** NHE9 mRNA is not significantly changed upon NHE6 KD in adipocytes. **(D)** NHE6 mRNA expression of adipocytes treated with shRNA (scramble or shNHE6) *post*-differentiation. **(E)** Representative confocal images showing reduction of GLUT4 at the cell surface of 3T3-L1 adipocytes following treatment of adipocytes with 100 nM insulin. Adipocytes were treated with shRNA (scramble or shNHE6) *post*-differentiation. Confocal images show a single plane through the midpoint of each cell. Images were taken at 63× magnification, (scale bar 10 µm). Graphs show mean ± SD for n=3 biological replicates per group. Data were evaluated using *t-*test. *\*\*P<0.01,* \*\*\**\*P<0.0001.* ns = not significant.

**Figure S2.**
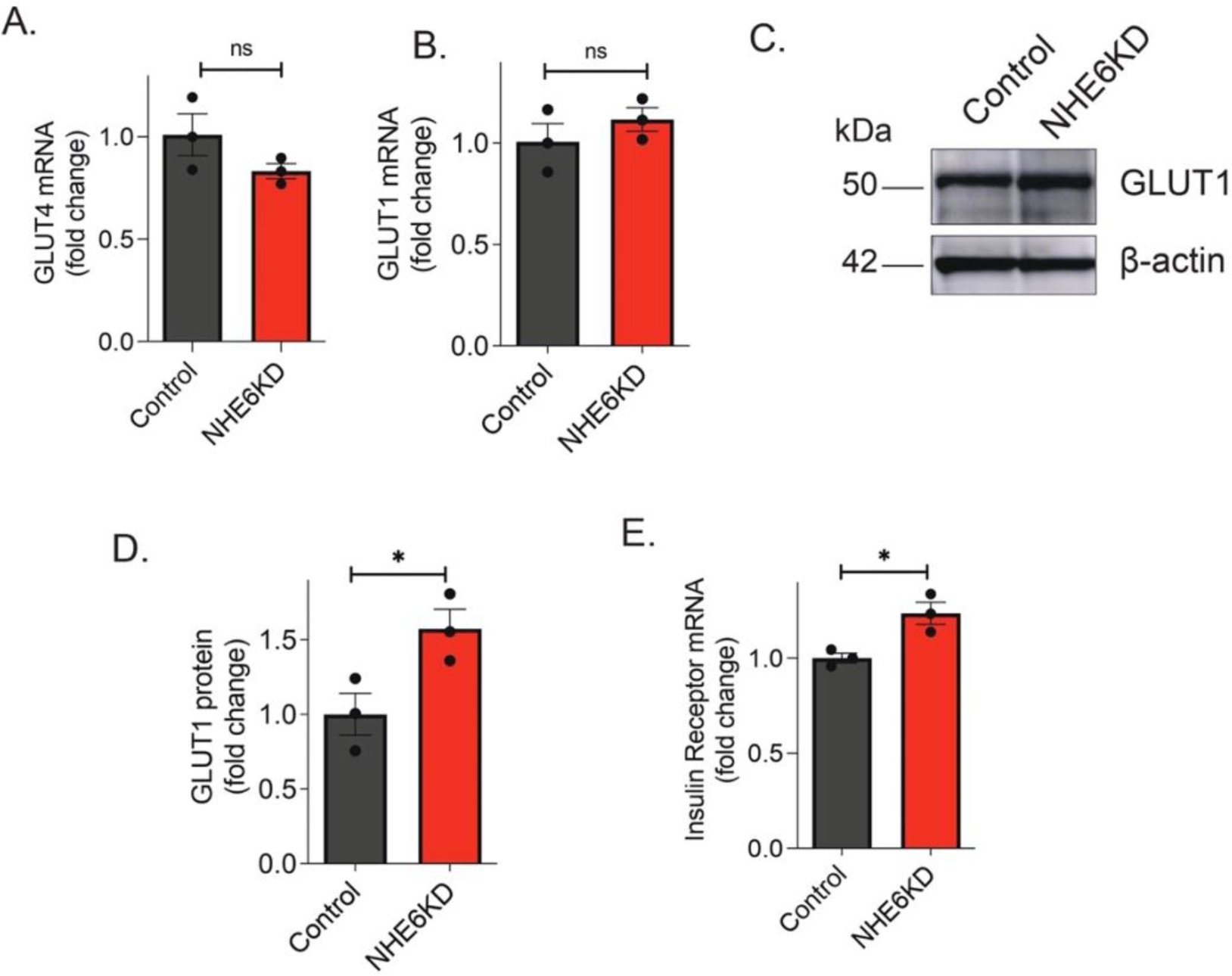
Effect of NHE6 knockdown on selected transcript and protein levels in adipocytes. **(A)** GLUT4 mRNA is not significantly altered due to NHE6 knockdown. **(B)** NHE6 knockdown does not change mRNA levels of GLUT1. **(C)** Representative Western blot of total protein levels of GLUT1 in control and NHE6 knockdown cells under basal conditions. **(D)** Densitometry of Western blot shows knockdown of NHE6 increases (0.57 +/-0.19) GLUT1 total protein. **(E)** Total mRNA of insulin receptor increases 24% upon NHE6 knockdown. Data were evaluated using *t-*test, n=3 independent experiments. *\*P<0.05,* \**\*P<0.01*.

**Figure S3.**
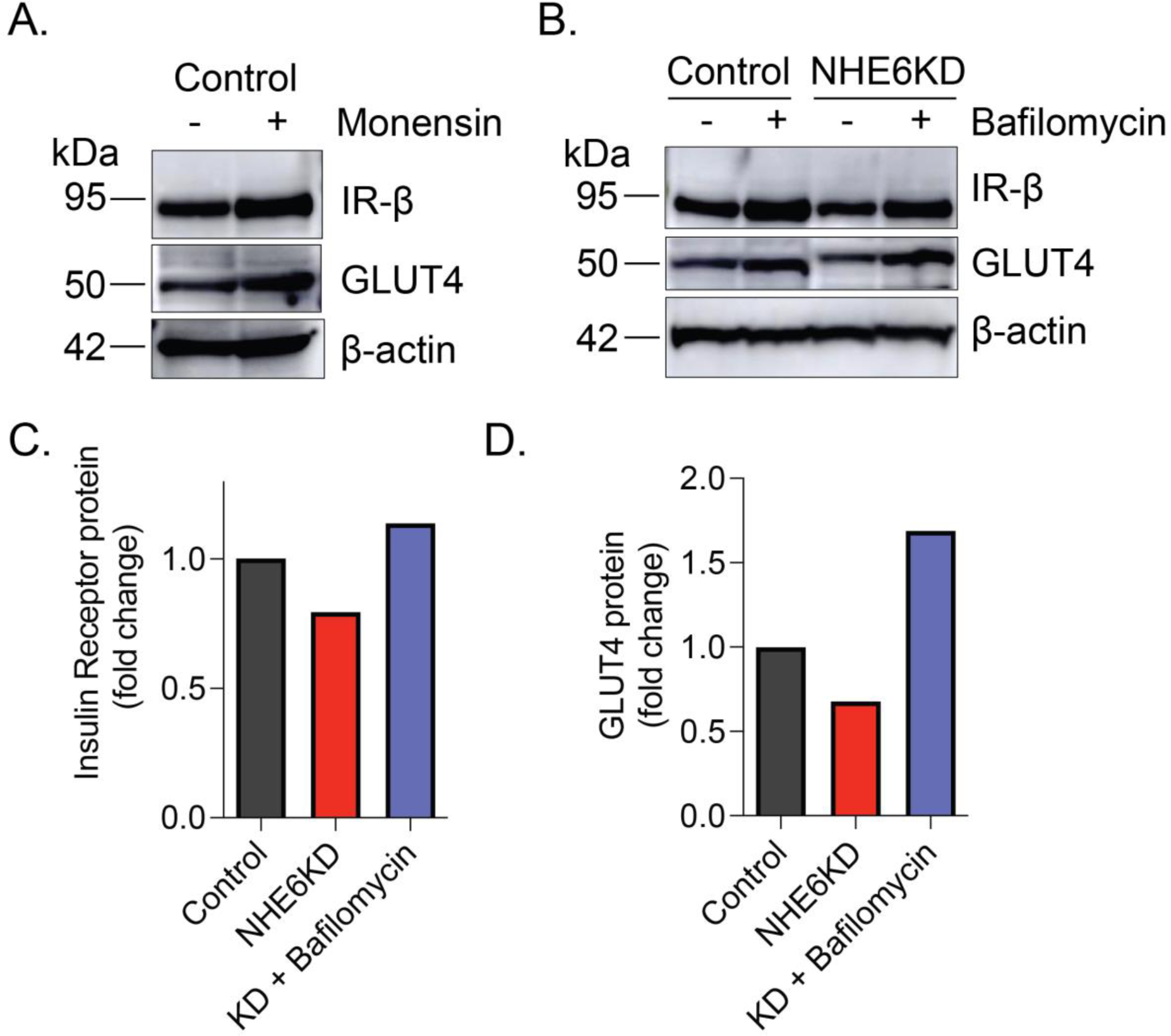
Chemical rescue of NHE6 KD in 3T3-L1 adipocytes with monensin and bafilomycin. **(A)** Western blot indicating treatment of control 3T3-L1 cMyc-GLUT4-mCherry adipocytes with 1 uM monensin for 16 hrs results in an increase in total protein of insulin receptor (IR-β) and GLUT4 in control adipocytes. **(B)** Western blot showing the effect of 25 nM bafilomycin treatment on total protein levels of both insulin receptor and GLUT4 in 3T3-L1 adipocytes. **(C-D)** Densitometry analysis of Western blot shown in **(B)** illustrating rescue of insulin receptor protein **(C)** and GLUT4 protein **(D)** in NHE6 KD following 2 hr treatment with bafilomycin. Graphs show quantitation of Western blot, n = 1.

## References

1. Bryant, N.J., R. Govers, and D.E. James, Regulated transport of the glucose transporter GLUT4. Nat Rev Mol Cell Biol, 2002. 3(4): p. 267–77.

2. Klip, A., T.E. McGraw, and D.E. James, Thirty sweet years of GLUT4. J Biol Chem, 2019. 294(30): p. 11369–11381.

3. Garvey, W.T., et al., Gene expression of GLUT4 in skeletal muscle from insulin-resistant patients with obesity, IGT, GDM, and NIDDM. Diabetes, 1992. 41(4): p. 465–75.

4. Kim, J. and Y.J. You, Regulation of organelle function by metformin. IUBMB Life, 2017. 69(7): p. 459–469.

5. Grisouard, J., et al., Mechanisms of metformin action on glucose transport and metabolism in human adipocytes. Biochemical Pharmacology, 2010. 80(11): p. 1736–1745.

6. Chinni, S.R. and A. Shisheva, Arrest of endosome acidification by bafilomycin A1 mimics insulin action on GLUT4 translocation in 3T3-L1 adipocytes. Biochem J, 1999. 339 (Pt 3)(Pt 3): p. 599–606.

7. Brett, C.L., M. Donowitz, and R. Rao, Evolutionary origins of eukaryotic sodium/proton exchangers. Am J Physiol Cell Physiol, 2005. 288(2): p. C223–39.

8. Kondapalli, K.C., et al., A leak pathway for luminal protons in endosomes drives oncogenic signalling in glioblastoma. Nat Commun, 2015. 6: p. 6289.

9. Ko, M., et al., The endosomal pH regulator NHE9 is a driver of stemness in glioblastoma. PNAS Nexus, 2022.

10. Ouyang, Q., et al., Christianson Syndrome Protein NHE6 Modulates TrkB Endosomal Signaling Required for Neuronal Circuit Development. Neuron, 2013.

11. Prasad, H. and R. Rao, Amyloid clearance defect in ApoE4 astrocytes is reversed by epigenetic correction of endosomal pH. Proc Natl Acad Sci U S A, 2018. 115(28): p. E6640–E6649.

12. Prasad, H., et al., NHA2 promotes cyst development in an in vitro model of polycystic kidney disease. J Physiol, 2019. 597(2): p. 499–519.

13. Kim, J., et al., NHX-5, an Endosomal Na+/H+ Exchanger, Is Associated with Metformin Action. J Biol Chem, 2016. 291(35): p. 18591–9.

14. Slack, C., A. Foley, and L. Partridge, Activation of AMPK by the putative dietary restriction mimetic metformin is insufficient to extend lifespan in Drosophila. PLoS One, 2012. 7(10): p. e47699.

15. Prasad, H. and R. Rao, Histone deacetylase-mediated regulation of endolysosomal pH. J Biol Chem, 2018. 293(18): p. 6721–6735.

16. Prior, M.J., et al., Quantitative proteomic analysis of the adipocyte plasma membrane. J Proteome Res, 2011. 10(11): p. 4970–82.

17. Zebisch, K., et al., Protocol for effective differentiation of 3T3-L1 cells to adipocytes. Anal Biochem, 2012. 425(1): p. 88–90.

18. Lim, C.Y., et al., Tropomodulin3 is a novel Akt2 effector regulating insulin-stimulated GLUT4 exocytosis through cortical actin remodeling. Nat Commun, 2015. 6: p. 5951.

19. Lorenzo, D.N. and V. Bennett, Cell-autonomous adiposity through increased cell surface GLUT4 due to ankyrin-B deficiency. Proc Natl Acad Sci U S A, 2017. 114(48): p. 12743–12748.

20. Takahashi, Y., et al., Perilipin2 plays a positive role in adipocytes during lipolysis by escaping proteasomal degradation. Sci Rep, 2016. 6: p. 20975.

21. Lee, C., et al., A two-domain elevator mechanism for sodium/proton antiport. Nature, 2013. 501(7468): p. 573–7.

22. Zhao, H., et al., Emerging roles of Na(+)/H(+) exchangers in epilepsy and developmental brain disorders. Prog Neurobiol, 2016. 138-140: p. 19–35.

23. Brett, C.L., et al., Human Na(+)/H(+) exchanger isoform 6 is found in recycling endosomes of cells, not in mitochondria. Am J Physiol Cell Physiol, 2002. 282(5): p. C1031–41.

24. Hill, J.K., et al., Vestibular hair bundles control pH with (Na+, K+)/H+ exchangers NHE6 and NHE9. J Neurosci, 2006. 26(39): p. 9944–55.

25. Piper, R.C., L.J. Hess, and D.E. James, Differential sorting of two glucose transporters expressed in insulin-sensitive cells. Am J Physiol, 1991. 260(3 Pt 1): p. C570–80.

26. Tanner, L.I. and G.E. Lienhard, Insulin elicits a redistribution of transferrin receptors in 3T3-L1 adipocytes through an increase in the rate constant for receptor externalization. J Biol Chem, 1987. 262(19): p. 8975–80.

27. Volchuk, A., et al., Cellubrevin is a resident protein of insulin-sensitive GLUT4 glucose transporter vesicles in 3T3-L1 adipocytes. J Biol Chem, 1995. 270(14): p. 8233–40.

28. Wardzala, L.J., et al., Potential mechanism of the stimulatory action of insulin on insulin-like growth factor II binding to the isolated rat adipose cell. Apparent redistribution of receptors cycling between a large intracellular pool and the plasma membrane. J Biol Chem, 1984. 259(13): p. 8378–83.

29. Larance, M., et al., Characterization of the role of the Rab GTPase-activating protein AS160 in insulin-regulated GLUT4 trafficking. J Biol Chem, 2005. 280(45): p. 37803–13.

30. Leto, D. and A.R. Saltiel, Regulation of glucose transport by insulin: traffic control of GLUT4. Nat Rev Mol Cell Biol, 2012. 13(6): p. 383–96.

31. Kohn, A.D., et al., Construction and characterization of a conditionally active version of the serine/threonine kinase Akt. J Biol Chem, 1998. 273(19): p. 11937–43.

32. Ng, Y., et al., Rapid activation of Akt2 is sufficient to stimulate GLUT4 translocation in 3T3-L1 adipocytes. Cell Metab, 2008. 7(4): p. 348–56.

33. Shi, J. and K.V. Kandror, Sortilin is essential and sufficient for the formation of Glut4 storage vesicles in 3T3-L1 adipocytes. Dev Cell, 2005. 9(1): p. 99–108.

34. Jedrychowski, M.P., et al., Proteomic analysis of GLUT4 storage vesicles reveals LRP1 to be an important vesicle component and target of insulin signaling. J Biol Chem, 2010. 285(1): p. 104–14.

35. Kondapalli, K.C., H. Prasad, and R. Rao, An Inside Job: How Endosomal Na+/H+ Exchangers Link to Autism and Neurological Disease. Frontiers in Cellular Neuroscience, 2014. 8.

36. Prasad, H. and R. Rao, The Na+/H+ exchanger NHE6 modulates endosomal pH to control processing of amyloid precursor protein in a cell culture model of Alzheimer disease. J Biol Chem, 2015. 290(9): p. 5311–27.

37. Muro, S., et al., Control of endothelial targeting and intracellular delivery of therapeutic enzymes by modulating the size and shape of ICAM-1-targeted carriers. Mol Ther, 2008. 16(8): p. 1450–8.

38. Thong, F.S., C.B. Dugani, and A. Klip, Turning signals on and off: GLUT4 traffic in the insulin-signaling highway. Physiology (Bethesda), 2005. 20: p. 271–84.

39. Cignarelli, A., et al., Insulin and Insulin Receptors in Adipose Tissue Development. Int J Mol Sci, 2019. 20(3).

40. Ashrafi, G., et al., GLUT4 Mobilization Supports Energetic Demands of Active Synapses. Neuron, 2017. 93(3): p. 606–615.e3.

41. Gilfillan, G.D., et al., SLC9A6 mutations cause X-linked mental retardation, microcephaly, epilepsy, and ataxia, a phenotype mimicking Angelman syndrome. Am J Hum Genet, 2008. 82(4): p. 1003–10.

42. Christianson, A.L., et al., X linked severe mental retardation, craniofacial dysmorphology, epilepsy, ophthalmoplegia, and cerebellar atrophy in a large South African kindred is localised to Xq24-q27. J Med Genet, 1999. 36(10): p. 759–66.

43. Schroer, R.J., et al., Natural history of Christianson syndrome. Am J Med Genet A, 2010. 152A(11): p. 2775–83.

44. Yamauchi, T., et al., The fat-derived hormone adiponectin reverses insulin resistance associated with both lipoatrophy and obesity. Nat Med, 2001. 7(8): p. 941–6.

45. Rossetti, L., et al., Peripheral but not hepatic insulin resistance in mice with one disrupted allele of the glucose transporter type 4 (GLUT4) gene. J Clin Invest, 1997. 100(7): p. 1831–9.

46. Abel, E.D., et al., Adipose-selective targeting of the GLUT4 gene impairs insulin action in muscle and liver. Nature, 2001. 409(6821): p. 729–33.

47. Stockli, J., D.J. Fazakerley, and D.E. James, GLUT4 exocytosis. J Cell Sci, 2011. 124(Pt 24): p. 4147–59.

